# A pivotal role of serine 225 phosphorylation in the function of hepatitis C virus NS5A revealed with the application of a phosphopeptide antiserum and super-resolution microscopy

**DOI:** 10.1101/387407

**Authors:** Niluka Goonawardane, Chunhong Yin, Mark Harris

## Abstract

NS5A is a multi-functional phosphoprotein that plays a key role in both viral replication and assembly. The identity of the kinases that phosphorylate NS5A, and the consequences for HCV biology, remain largely undefined. We previously identified serine 225 (S225) within low complexity sequence (LCS) I as a major phosphorylation site and used a phosphoablatant mutant (S225A) to define a role for S225 phosphorylation in the regulation of genome replication, interactions of NS5A with several host proteins and the sub-cellular localisation of NS5A. To investigate this further, we raised an antiserum to S225 phosphorylated NS5A (pS225). Western blot analysis revealed that pS225 was exclusively found in the hyper-phosphorylated NS5A species. Furthermore, using kinase inhibitors we demonstrated that S225 was phosphorylated by casein kinase 1α (CK1α) and polo-like kinase 1 (PLK1). Using a panel of phosphoablatant mutants of other phosphorylation sites in LCSI we obtained the first direct evidence of bidirectional hierarchical phosphorylation initiated by phosphorylation at S225.

Using super-resolution microscopy (Airyscan and Expansion), we revealed a unique architecture of NS5A-positive clusters in HCV-infected cells - pS225 was concentrated on the surface of these clusters, close to lipid droplets. Pharmacological inhibition of S225 phosphorylation resulted in the condensation of NS5A-positive clusters into larger structures, recapitulating the S225A phenotype. Although S225 phosphorylation was not specifically affected by daclatasvir treatment, the latter also resulted in a similar condensation. These data are consistent with a key role for S225 phosphorylation in the regulation of NS5A function.

**Importance:** NS5A has obligatory roles in the hepatitis C virus lifecycle, and is proposed to be regulated by phosphorylation. As NS5A is a target for highly effective direct-acting antivirals (DAAs) such as daclatasvir (DCV) it is vital to understand how phosphorylation occurs and regulates NS5A function. We previously identified serine 225 (S225) as a major phosphorylation site. Here we used an antiserum specific for NS5A phosphorylated at S225 (pS225-NS5A) to identify which kinases phosphorylate this residue. Using super-resolution microscopy we showed that pS225 was present in foci on the surface of larger NS5A-positive clusters likely representing genome replication complexes. This location would enable pS225-NS5A to interact with cellular proteins and regulate the function and distribution of these complexes. Both loss of pS225 and DCV treatment resulted in similar changes to the structure of these complexes, suggesting that DAA treatment might target a function of NS5A that is also regulated by phosphorylation.

## Introduction

Hepatitis C virus (HCV) infects an estimated 73 million people and frequently results in chronic liver disease – cirrhosis and hepatocellular carcinoma (1-3). HCV, a member of the *Hepacivirus* genus within the *Flaviviridae* virus family, has a 9.6 kb positive-stranded RNA genome with 5′ and 3′ untranslated regions (UTRs) flanking a single open reading frame encoding a 3,000-residue polyprotein precursor. This is co- and post-translationally processed by host and viral proteases to yield four structural proteins (core, E1, E2, and p7) and six non-structural proteins (NS2, NS3, NS4A, NS4B, NS5A, and NS5B) (4-6). The NS proteins play a number of roles during the HCV life cycle (6). NS3-NS5B are necessary and sufficient for HCV genome replication in association with multiple-membrane vesicles (MMV) (7), whose formation is induced by NS4B and NS5A. MMVs compartmentalise genome replication complexes (RC) (8-10) shielding them from host defence mechanisms and concentrating components to stimulate replicative efficiency. Polyprotein processing and genome replication are the targets for recently developed direct-acting antivirals (DAAs), used in combination therapies to effectively treat HCV infection. DAAs target NS3/4A protease, NS5A, and NS5B RNA-dependent RNA polymerase, revolutionising HCV therapy (11). Daclatasvir (DCV) exemplifies a class of DAAs proposed to be potent inhibitors of NS5A (12, 13). However, the mode(s) of action for these compounds remains obscure as it is not clear which of the myriad functions of NS5A are inhibited.

NS5A is a multifunctional protein comprising three domains (DI-III) and an N-terminal amphipathic helix anchoring it to cytoplasmic membranes (Fig 1A). The structure of DI has been determined (14-16), in contrast DII and DIII are intrinsically disordered, with elements of secondary structure (17). The domains are linked by low complexity sequences (LCS) I and II (Fig 1A), LCSI is serine-rich, and LCSII is proline-rich. All three domains bind the HCV 3′UTR (18): DI functions in both genome replication and virus assembly (19), DII is exclusively involved in genome replication (20) and DIII plays a critical role in virion assembly (21). NS5A has been shown to associate with lipid droplets (LDs) in infected cells (22), where it is postulated to physically link the processes of genome replication and virus assembly (19). NS5A also interacts with many cellular proteins (23, 24) and signalling pathways (25-27), modifying the host cell environment to favour the virus, this may require a subset of NS5A distinct from that involved in genome replication or virus assembly.

**Figure 1.**
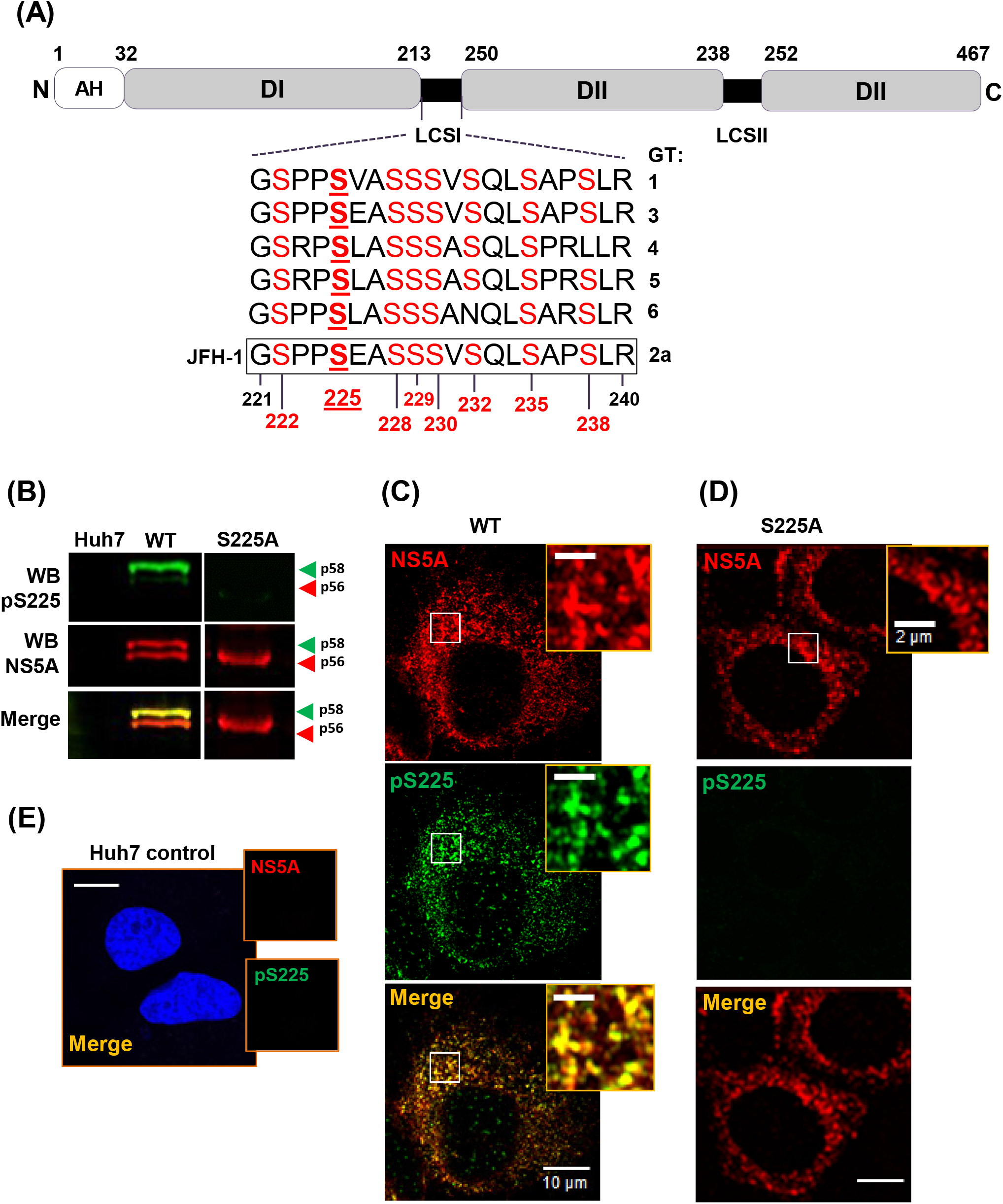
Validation of pS225 antiserum. (A) Schematic representation of the domain organization of NS5A. The three domains (I-III), the linking low complexity sequences (LCSI and II), and the membrane anchoring amphipathic helix (AH) are illustrated as are the peptide sequences of the LCSI region from six major HCV genotypes. Numbers indicate amino acid positions in the JFH-1 genotype 2a NS5A sequence. (B) Specificity of the pS225 antibody. Lysates from Huh7 cells electroporated with either wildtype (WT) or S225A mJFH-1 RNA were analyzed by Western blot with either rabbit-anti-pS225 (green), or sheep-anti-NS5A (red) polyclonal antisera. (C-E) Huh7 cells electroporated with WT (C) or S225A (D) mJFH-1 RNA were analysed by immunofluorescence with anti-pS225 or anti-NS5A antisera. (E) Uninfected Huh7 cells were used as negative control. Scale bars: 10 μm and 2 μm (insets).

Recently, NS5A phosphorylation has been extensively investigated to address how this post-translational modification might modulate the various functions of NS5A, protein conformation and protein-protein interactions. NS5A exists as two species with different mobility on SDS-PAGE termed basally-phosphorylated (apparent molecular weight 56kDa), and hyper-phosphorylated (58kDa). Of note the cell culture infectious JFH-1 (genotype 2a) HCV isolate used here, and most other studies, contains an 18-residue insertion in DIII compared to genotype 1b (14), thus the two JFH-1 species migrate with apparent molecular weights of 63 and 65kDa, however for consistency with the literature they will be referred to as p56 and p58. NS5A is phosphorylated on multiple serine and (to a lesser extent) threonine residues (28). Whereas p56 is reportedly phosphorylated within DII and DIII, in contrast, multiple phosphorylation events within the serine-rich LCSI result in the production of p58 (Fig 1A) (23, 29-31).

Two key questions regarding NS5A phosphorylation remain to be unambiguously answered: namely, which cellular kinases are responsible for phosphorylation and whether there is an order, or hierarchy, such that a priming phosphorylation event enables subsequent phosphorylation? In this regard LCSI is reminiscent of cellular proteins that exhibit hierarchical phosphorylation, for example β-catenin, in which serine 45 (S45) is phosphorylated by casein kinase I (CKI), priming it for phosphorylation at S41/37/33 by glycogen synthase kinase-3 (GSK-3) (32), triggering ubiquitinylation and proteosomal degradation of β-catenin. Intriguingly, CKIα has been implicated in production of the NS5A p58 as indicated by the loss of p58 following treatment of infected cells with CKIα-inhibitors (33, 34). A number of studies have used biochemical and genetic approaches to demonstrate that CKIα phosphorylates S225, S232, S235 and S238 but the relative importance of these phosphorylation events and their sequential order remains to be determined. Additionally, CKIα-phosphorylation requires priming phosphorylation at the −3 position, suggesting that other kinases may be involved (35). Indeed, another serine-threonine kinase, Polo-like kinase 1(PLK1), and the lipid kinase, phosphatidylinositol-4 kinase IIIα (PI4KIIIα) have also been shown to play a role in p58 production (36).

Our previous studies had pointed to a key role for S225 phosphorylation – the S225A phosphoablatant mutant had a dramatic phenotype: a loss of p58, a 10-fold reduction in JFH-1 genome replication, a lack of interactions between NS5A and a range of host proteins, and a perinuclear restricted localisation of NS5A and other replication complex components (23, 37). This phenotype prompted us to investigate the role of S225 phosphorylation more directly, therefore to complement the analysis of S225 mutants, we raised an antiserum to pS225-NS5A and used this unique reagent to probe the functions of S225-phosphorylated wildtype NS5A. We demonstrate that pS225 is exclusively present in p58 and results from the activity of both CK1α and PLK1. We also provide evidence that pS225 is the ‘priming’ phosphorylation event that drives subsequent phosphorylation across LCSI. Using super-resolution microscopy approaches we also reveal the detailed architecture of NS5A-positive clusters whereby in HCV-infected cells pS225 is not uniformly distributed, but is instead only present on the surface and in ‘foci’. Super-resolution analysis also revealed that treatment with DCV results in profound disruption to the architecture of these clusters – they become larger, less ordered and more diffuse. The same morphology was observed in the context of the S225A phosphoablatant mutant. The combination of the pS225 antiserum and super-resolution microscopy thus revealed unique insights into the role of NS5A phosphorylation and are consistent with a key role for S225 phosphorylation in the regulation of NS5A functions during the HCV lifecycle.

## Results

### Characterisation of an antiserum specific for S225 phosphorylated NS5A

We previously demonstrated a role for S225 phosphorylation in HCV genome replication, and further showed that this was likely mediated via interactions of NS5A with a defined subset of cellular factors (23, 37). A caveat to these studies was the necessary dependence on interpretation of data based on the analysis of the phenotype of S225 mutants (S225A phosphoablatant or S225D phosphomimetic). To address this we raised a rabbit polyclonal antiserum specific for the S225 phosphorylated form of NS5A using an appropriate phosphopeptide as immunogen (CARGSPP**pS**EASSS). The resulting anti-pS225 antiserum was validated by immunoblotting (Fig 1B) and reassuringly detected a single NS5A band in a lysate from Huh7 cells infected with wildtype HCV (mJFH-1). Consistent with the previous observation that the S225A phosphoablatant mutant resulted in a loss of p58 (37), the pS225 reactive species corresponded with this form of NS5A, illustrated by the merged image of pS225 (green) and total NS5A (red) signals. As expected, the pS225 antiserum exhibited no reactivity against the S225A mutant.

We also validated the use of this antiserum for immunofluorescence analysis by confocal microscopy. Huh7 cells stably harbouring a wildtype SGR were stained for total NS5A (red) and pS225 (green) (Fig 1C). The results were consistent with the western blot analysis, showing that in wildtype SGR-harbouring cells pS225 distribution significantly co-localised with the total NS5A staining. Even though the western blot data showed that a proportion of NS5A (p56) did not contain pS225, we expected to observe a uniform distribution of pS225 coincident with the total NS5A distribution. However, this was not the case – the co-localisation was not complete and there was a subset of total NS5A reactivity that did not stain for pS225. This observation is developed further later in this manuscript (Figs 7, 9). The specificity of the pS225 antiserum was further confirmed by a lack of cross-reactivity with either the S225A phosphoablatant NS5A or control cells (Fig 1D, E). Taken together these data validated the subsequent use of this unique reagent as a tool to investigate the properties of S225-phosphorylated NS5A in more detail.

### Identification of kinases that phosphorylate S225

We proceeded to use the antiserum to investigate which cellular kinases might be responsible for phosphorylation of S225. Previous studies (29, 33, 38), coupled with bioinformatic analysis (39), suggested that S225 was a potential phosphorylation site for three kinases: casein kinase-Iα (CKIα), polo-like kinase-1 (PLK1) and glycogen synthase kinase-3β (GSK3β). To investigate their potential roles in phosphorylation of S225, we treated HCV-infected Huh7 cells with three selective kinase inhibitors: D4476 (CKIα inhibitor); SBE13-HCl (PLK1 inhibitor), or GSK-3β Inhibitor VIII (GSK3β inhibitor). Huh7 cells were electroporated with mJFH-1 RNA, incubated for 24 h, and treated with kinase inhibitors for a further 24 h prior to harvest and analysis of lysates by western blot for total NS5A and pS225-NS5A (Fig 2A, B). The effects of the inhibitors on HCV genome replication were also assessed in parallel using Huh7 cells stably harbouring an HCV SGR expressing firefly luciferase-neomycin phosphotransferase (Feo) (Fig 2C). Cell viability was monitored by MTT assay (Fig 2D).

**Figure 2.**
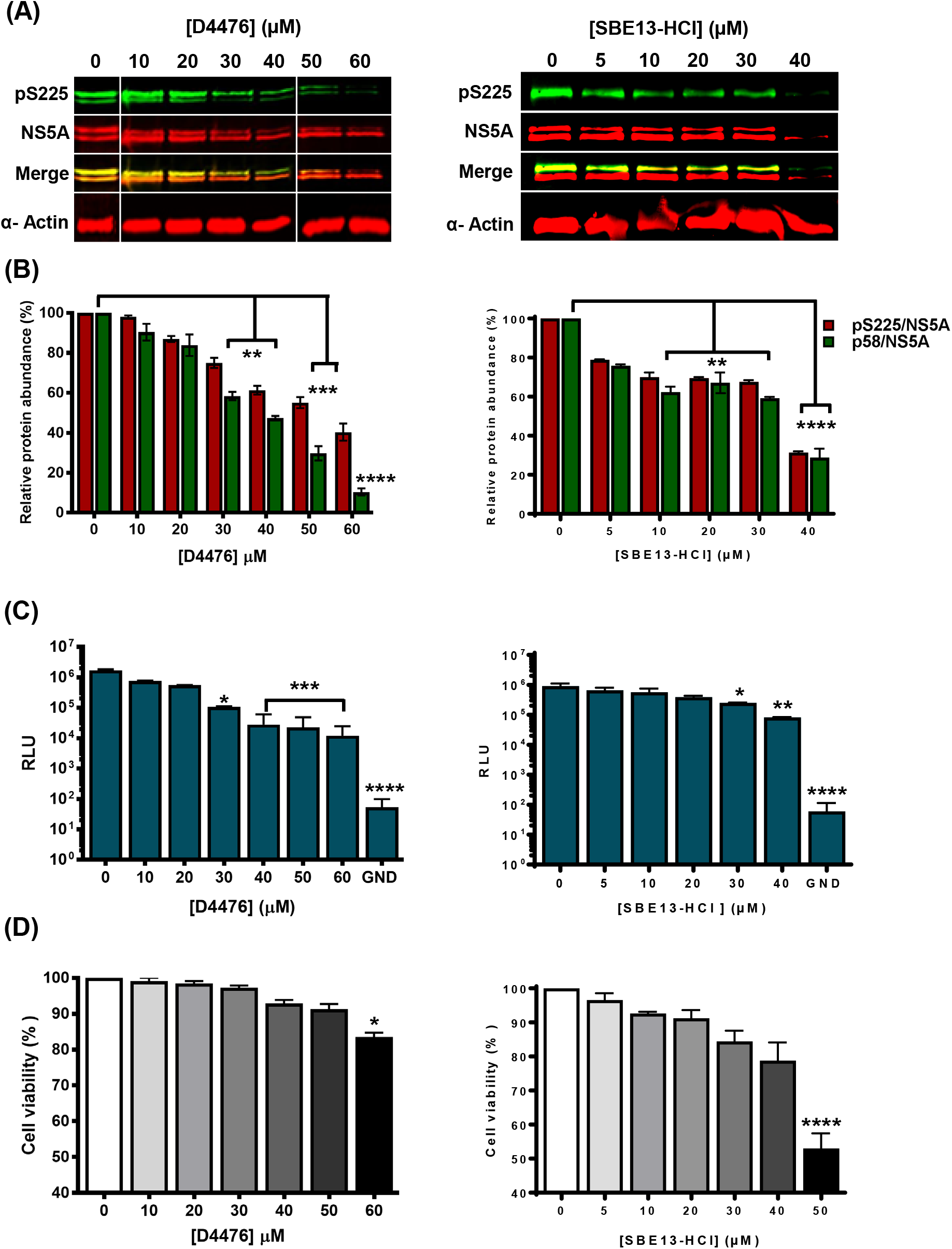
CK1α and PLK1 inhibition reduced NS5A S225 phosphorylation. (A) Huh7 cells were electroporated with WT mJFH-1 RNA and treated with D4476 or SBE13-HCl for 24 h prior to lysis and western blot analysis for pS225, total NS5A and α-actin. (B) Relative protein abundance for three independent experiments. Values are mean ± SD, normalized against values of the vehicle control (0 μM). (C) Assay for replication of SGR-Feo-JFH-1 at 24 h post inhibitor treatment. (D) Cell viability tested following 24 h inhibitor treatment assessed by MTT assay. Statistical analysis by paired t-tests: * P<0.05, ** P<0.01, *** P<0.001, **** P>0.0001.

Both D4476 and SBE13-HCl significantly reduced pS225 and p58 abundance in a dose-dependent manner (Fig 2A). Quantification of multiple western blots revealed that at the maximal non-cytotoxic concentrations (D4476: 50 μM, SBE13-HCl: 40 μM) phosphorylation of S225 was reduced by approximately 70% (Fig 2B). Consistent with the reduction in both pS225 and p58 abundance, D4476 or SBE13-HCl at the maximal non-cytotoxic concentrations also resulted in a 75-fold or 11-fold reduction in RNA replication respectively (Fig 2C). In contrast to the inhibitors of CKIα and PLK1, treatment with GSK-3β Inhibitor VIII had no effect on pS225 phosphorylation or HCV RNA replication (Fig S1). We conclude that S225 is a substrate for phosphorylation by either of the two cellular kinases, CKIα and PLK1, but is not phosphorylated by GSK-3β.

In parallel inhibitor-treated HCV-infected cells were analysed using high resolution confocal microscopy (Airyscan) to examine effects on the subcellular distribution of total NS5A, pS225-NS5A and LDs. The latter were chosen as we, and others, had previously shown co-localisation of NS5A and LD (19, 22, 30, 37, 40, 41), and redistribution of LDs in the context of cells infected with HCV containing the phosphoablatant S225A NS5A mutant (37). Treatment with D4476 resulted in a dose-dependent condensation of NS5A-positive punctae into larger structures, some of which accumulated at the surface of LD, particularly at higher concentrations (Fig 3A). Despite this accumulation, quantification of fluorescence images revealed that inhibitor treatment led to a reduction in the total amount of pS225-NS5A localised with LDs. (Fig 3B).

**Figure 3.**
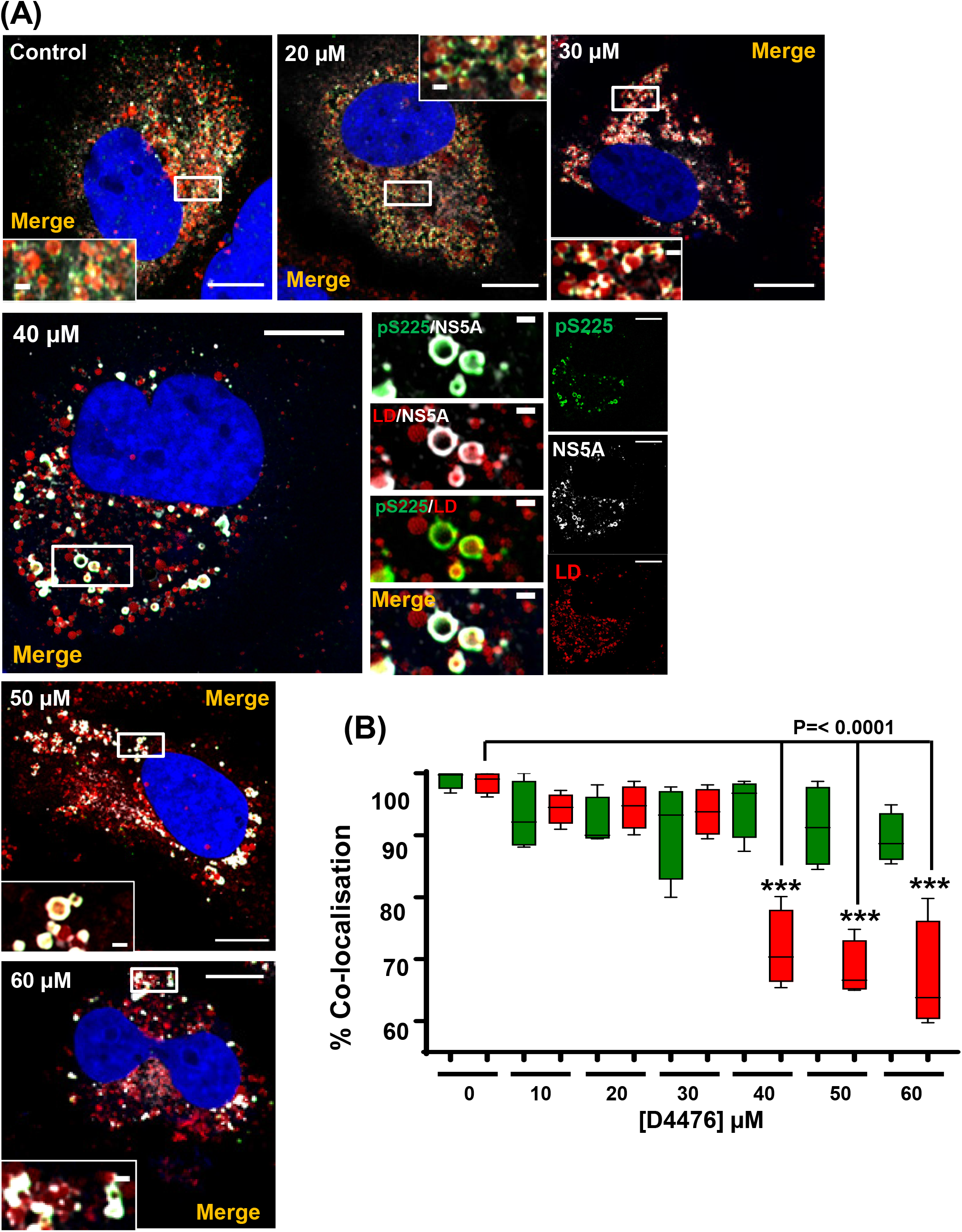
Effect of CK1a inhibition on the subcellular distribution of total NS5A, pS225-NS5A, and LD. (A) Immunofluorescence (IF) analysis for pS225-NS5A, total NS5A and LD. Huh7 cells were electroporated with WT mJFH-1 RNA and treated with the indicated concentration of the CKIα inhibitor D4476 from 24-48 h.p.e. Cells were fixed and stained for total NS5A (white), pS225 (green) and LD (red), prior to imaging by Airyscan microscopy. Scale bars: 10 μm and 1 μm, respectively. (B) Quantification of the percentages of pS225-NS5A colocalised with LD (green), or LD colocalised with pS225-NS5A (red) from 10 cells for each inhibitor concentration. **** P<0.0001, compared to control (0 μM) data.

In comparison, treatment with SBE13-HCl had a less dramatic effect on the distribution of pS225-NS5A (Fig. 4A), although a degree of LD accumulation was observed. Consistent with this, quantification showed a small but significant reduction in the amount of pS225-NS5A localised with LDs at high SBE13-HCl concentrations (Fig 5B). Although inhibition of GSK3β did not affect levels of pS225-NS5A, RNA replication (Fig S1) or co-localisation between pS225-NS5A and LDs (Fig S2), it did result in condensation of NS5A-positive punctae into larger structures and a perinuclear concentration of LD (Fig S2A). This may be due to the role of GSK3β in lipid metabolism (42).

**Figure 4.**
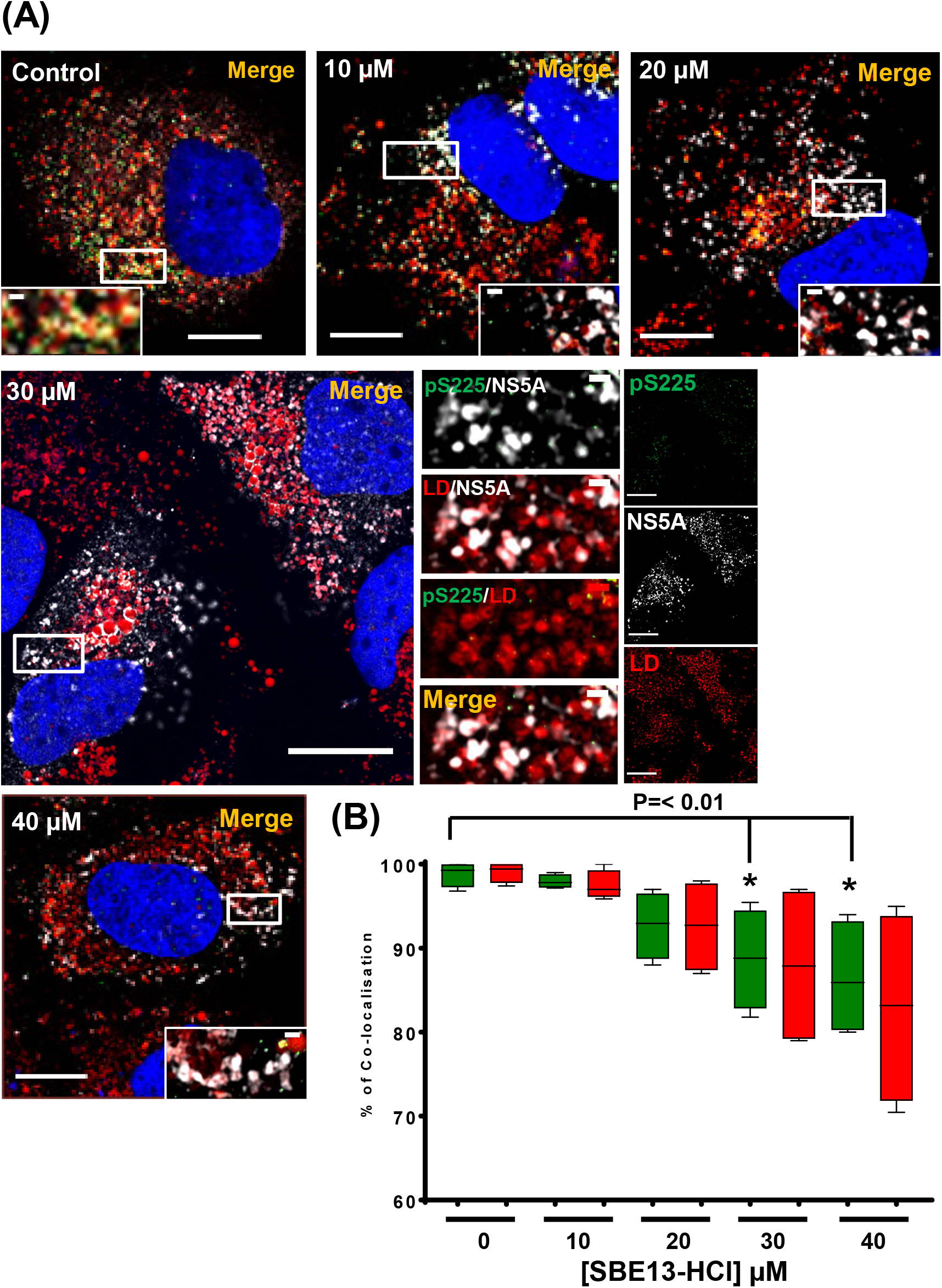
Effect of PLK inhibition on the subcellular distribution of total NS5A, pS225-NS5A, and LD. (A) Immunofluorescence (IF) analysis for pS225-NS5A, total NS5A and LD. Huh7 cells were electroporated with WT mJFH-1 RNA and treated with the indicated concentration of the PLK inhibitor SBE13-HCl from 24-48 h.p.e. Cells were fixed and stained for total NS5A (white), pS225 (green) and LD (red), prior to imaging by Airyscan microscopy. Scale bars: 10 μm and 1 μm, respectively. (B) Quantification of the percentages of pS225-NS5A colocalised with LD (green), or LD colocalised with pS225-NS5A (red) from 10 cells for each inhibitor concentration. * P<0.01 compared to control (0 μM) data.

**Figure 5.**
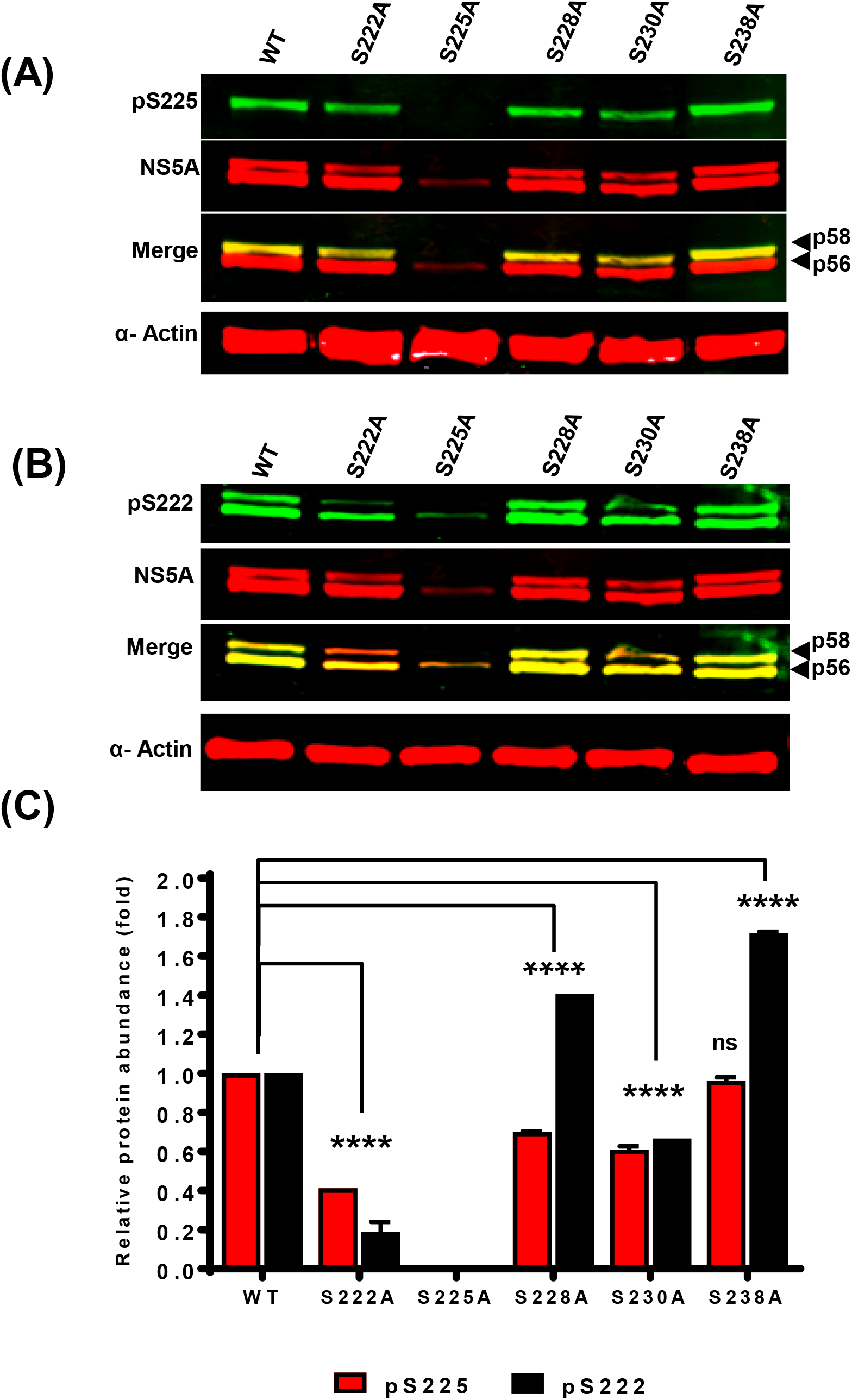
pS225 primes hierarchical phosphorylation. Huh7 cells were electroporated with the indicated mJFH-1 WT and mutant RNAs. Lysates were analysed by western blot for pS225 (A) or pS222 (B) (43), total NS5A and β-actin. (C) Quantification of pS225 (red) and pS222 (black) abundance from multiple western blots (*n*=3). Values are mean ± SE. **** P>0.0001.

### Evidence for a role of S225 phosphorylation in initiating hierarchical phosphorylation in LCSI

A number of lines of evidence points to a hierarchical, or sequential, phosphorylation cascade across LCSI. These include the conservation and spacing of the serine residues, and mass spectrometric evidence for multiply-phosphorylated species corresponding to LCSI (31, 38, 43). Additionally the group of Yu (33) recently generated pS235 and pS238-specific antisera and demonstrated that pS238 was dependent on pS235, but not vice-versa, suggesting that S235 was the primary phosphorylation site leading to a hierarchical phosphorylation of other serines within LCSI. However, they did not address the potential role of S225phosphorylation, and we hypothesised that this might precede phosphorylation of S235 and initiate the hierarchical phosphorylation cascade.

To provide evidence for this hypothesis we used western blot analysis to examine the presence of pS225 reactivity in lysates of cells infected with mJFH-1 phosphoablatant mutants: S222A, S225A, S228A, S230A and S238A. We were unable to interrogate the remaining phosphoablatant mutants as they were replication-inactive. As shown in Fig 5A, the only mutant which affected pS225 reactivity was S225A itself, indicating that this phosphorylation event was independent of other phosphorylated serines in LCSI.

In a previous study (43) we generated an antibody specific to pS222. Although this reagent was in scarce supply, we had sufficient to interrogate the same samples for the presence of pS222. As expected, S222A exhibited significantly reduced levels of pS222 within the p58 species (Fig 5B), however this mutant retained wildtype level of reactivity for the p56 species. We believe this reflects reactivity against the non-phosphorylated sequence and is due to the fact that this serum was not affinity purified using the phospho-peptide (as had the pS225 antiserum). Importantly, the only other phosphoablatant mutant that blocked pS222 was S225A. These observations are consistent with the suggestion that S225 phosphorylation is the ‘priming’ event that leads to a bi-directional hierarchical cascade of phosphorylation events.

We also used confocal microscopy with Airyscan to analyse the distribution of both total NS5A and pS225-NS5A in cells infected with the various phosphoablatant mutants. As shown in Fig 6, all mutants displayed a broad cytoplasmic distribution of total NS5A that was similar to wildtype, with the exception of S225A which exhibited a more compact perinuclear distribution, as documented previously (23). However, as observed in Fig 1, we noted again that in all cases there was a subset of total NS5A reactivity that did not stain for pS225, and this led us to investigate this observation in more detail. We thus undertook a temporal analysis of the distribution of total NS5A, pS225-NS5A and LDs in cells electroporated with mJFH-1 RNA. As shown in Fig 7, there was a noticeable increase in the amount of pS225 reactivity between 24-72 h.p.e. As observed previously (19), over time the size of LDs increased and NS5A became almost exclusively associated with the periphery of these LDs. The distribution of pS225 also associated with the periphery of LDs but was restricted to part of the LDs. For example, at 72 h.p.e. the zoomed image shows an LD where NS5A is associated with the majority of the periphery, but pS225 is restricted to distinct locations.

**Figure 6.**
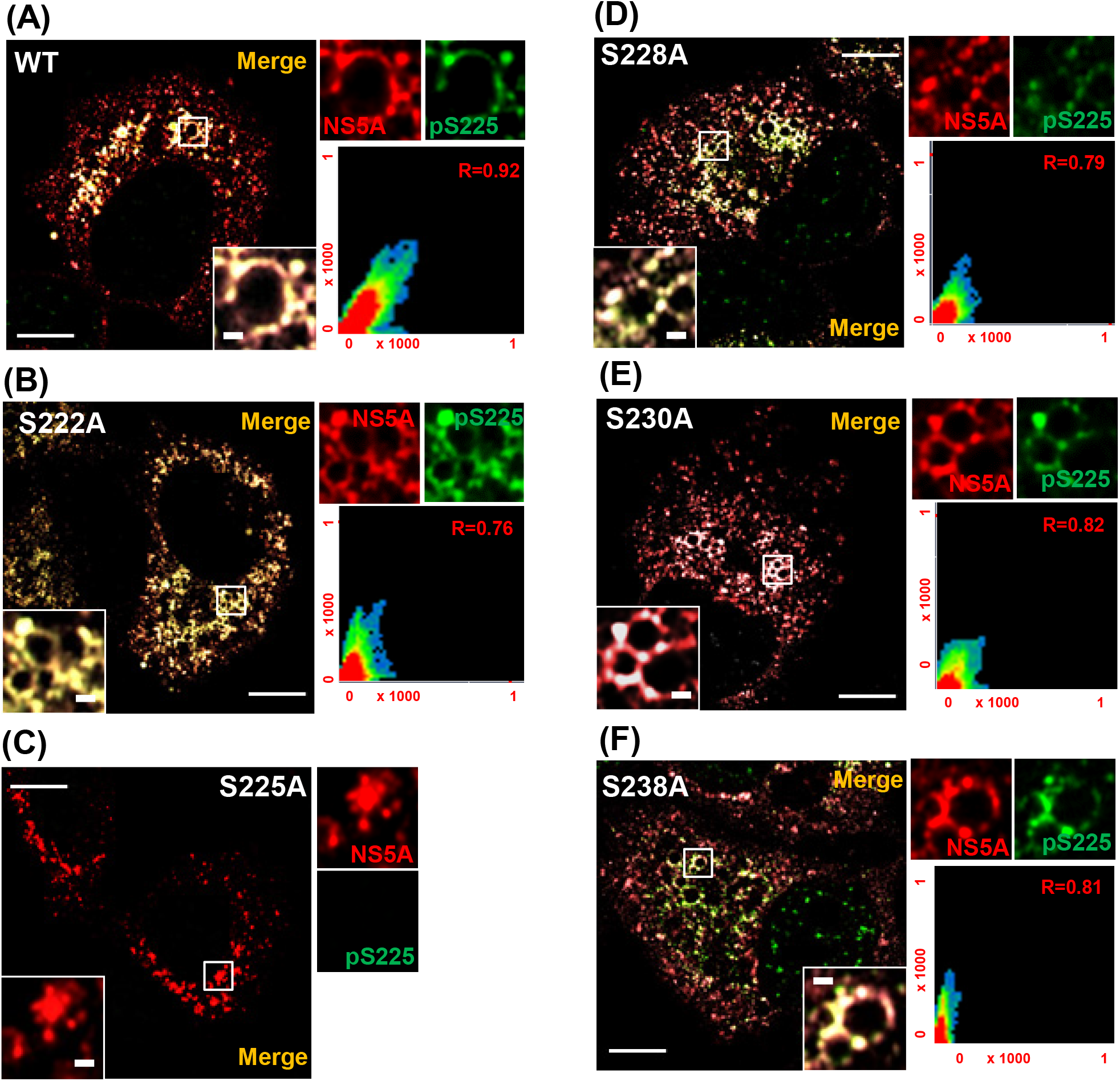
Subcellular distribution of phosphoablatant mutants of NS5A. Huh7 cells were electroporated with the indicated mJFH-1 WT and mutant RNAs. Cells were fixed and stained for total NS5A (red) and pS225 (green), prior to imaging by Airyscan microscopy. Scale bars: 10 μm and 1 μm, respectively. The scatter plots show the co-localization of NS5A and LD (*x* and *y* axes, pixel numbers). The correlation value (*r*) is shown for each image.

**Figure 7.**
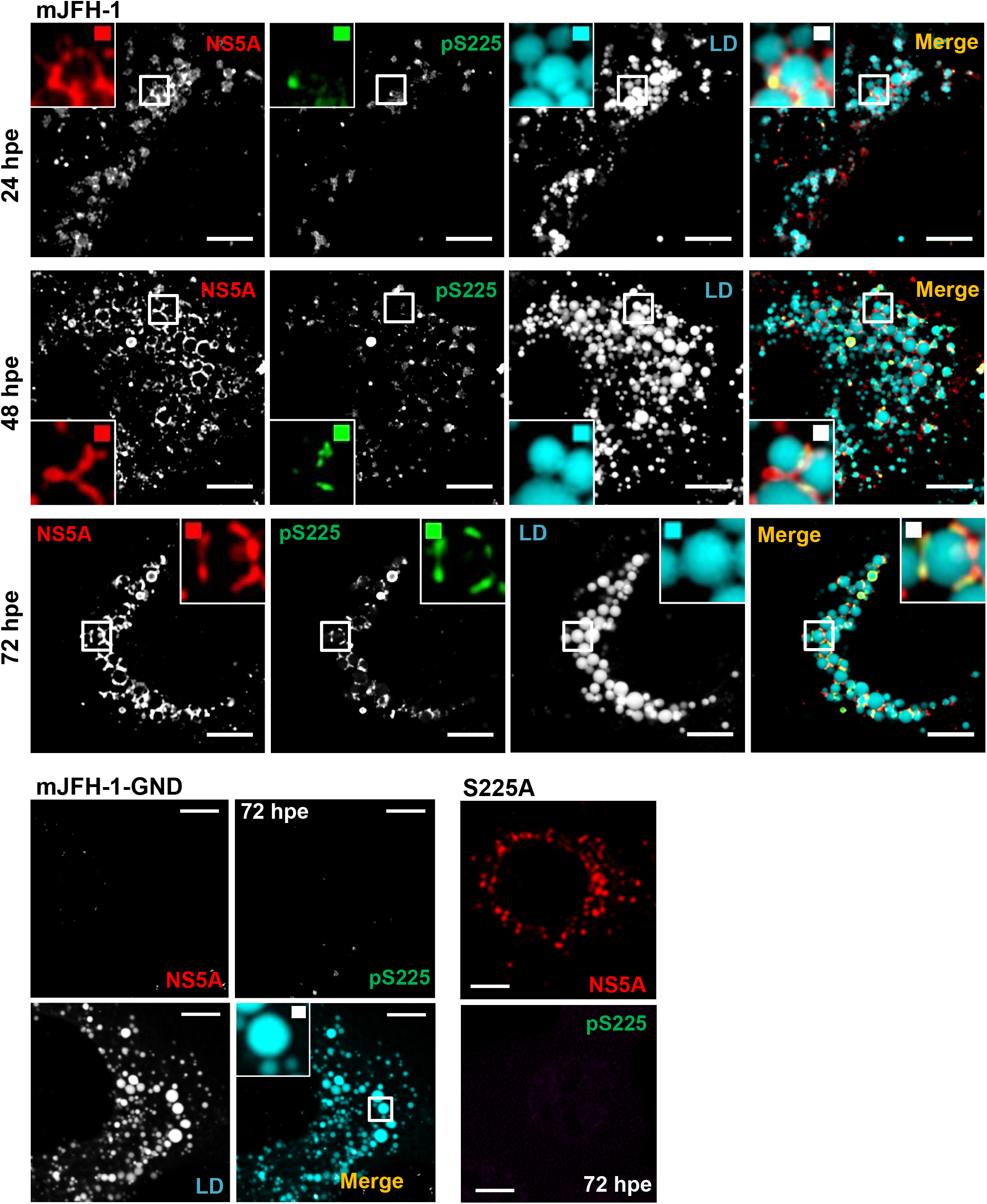
Pre-expansion microscopic analysis of NS5A and pS225 subcellular distribution in HCV infected cells. Huh7 cells were electroporated with either WT, S225A or GND (NS5B polymerase defective) mJFH-1 RNA, seeded onto coverslips and incubated for 24, 48 and 72 h. Cells were fixed and stained for total NS5A (red), pS225 (green) and LDs (Cyan) prior to imaging by confocal microscopy. Scale bars are 5 μm and 0.5 μm (insets) respectively.

### Expansion microscopy (ExM) imaging of NS5A and pS225 in HCV infected cells

Previously we had shown using direct stochastic optical reconstruction microscopy (dSTORM) that the S225A phosphoablatant mutant NS5A was present in larger cytoplasmic punctae in comparison to wildtype or the phosphomimetic S225D NS5A (23). In our laboratory at present dSTORM analysis is restricted to a single-colour, however, to assess the distribution of pS225 in comparison to total NS5A would require multi-colour capability. We therefore decided to utilise the novel super-resolution technique of expansion microscopy (ExM) (44-46). Unlike other super-resolution approaches which rely on specialised equipment and complex approaches to image analysis and interpretation, ExM relies on the simple principle of physical expansion of the sample. Briefly, cell samples are probed with fluorescently labelled antibodies and then embedded in a swellable gel matrix which chemically anchors the fluorescent labels. Samples are then physically expanded to ~4 times the original dimensions by swelling the gel in an aqueous environment. ExM thus effectively improves the diffraction-limited resolution by a factor of 4, achieving super-resolution imaging (~70-65 nm resolution) without needing to modify existing microscope hardware.

We first confirmed the capability of ExM to improve the resolution using our facilities (Zeiss LSM880). To do this we analysed vimentin intermediate filaments in Huh7 cells by immunofluorescence, and then compared the microscope images pre- and post-expansion (Fig 8). Whereas the pre-expansion image showed individual vimentin filaments with blurred edges (Fig 8A), the ExM image allowed clear distinction of the substructures within the filaments (Fig 8B). Quantification of the intensity profiles revealed more precise resolution of this sub-filamentous structure (Fig 8C). We therefore applied this technique to interrogate pS225-NS5A and total NS5A distribution in Huh7 cells electroporated with mJFH-1 RNA at different times.

**Figure 8.**
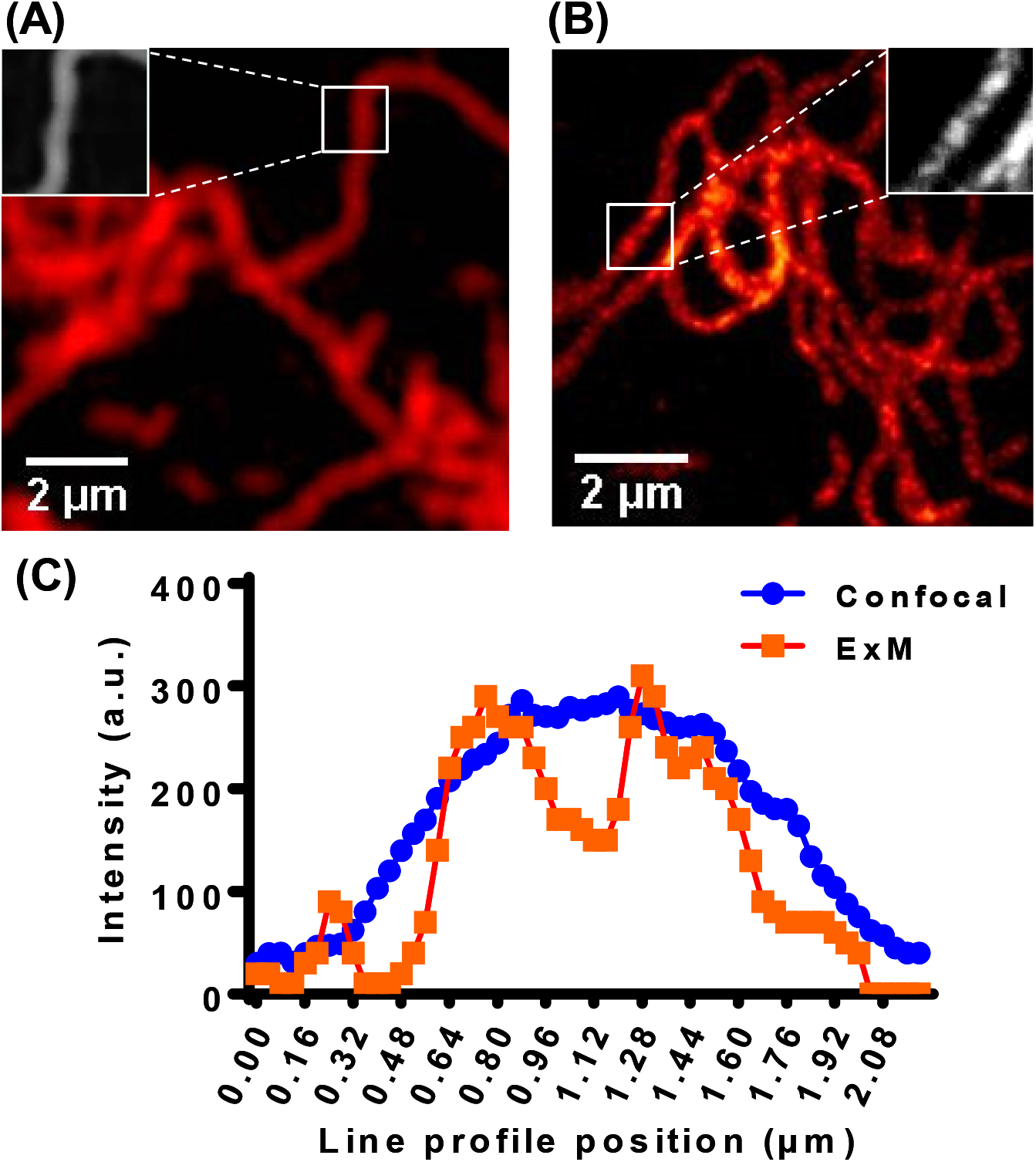
Validation of expansion microscopy (ExM) Huh7 cells were fixed and stained with a mouse polyclonal anti-vimentin antibody and secondary Alexa Fluor 488 antibody, and processed for ExM. **(A)** Pre-expansion image. **(B)** Post-expansion image. Magnified views of the selected regions are indicated. **(C)** Intensity profiles of images in magnified views.

As observed using Airyscan, ExM analysis revealed a diffuse distribution of NS5A throughout the cytoplasm (Fig 9A) with significant co-localisation between NS5A and pS225 (merge panel). Over time, the diffuse distribution resolved into a more clustered appearance with an overall increase in the size of NS5A-positive structures. Quantification confirmed this as the peak area of these structures increased from 0.09-0.12 μm^2^ at 24 h.p.e., to 0.15–0.18 μm^2^ at 72 h.p.e. (Fig 8B).

**Figure 9.**
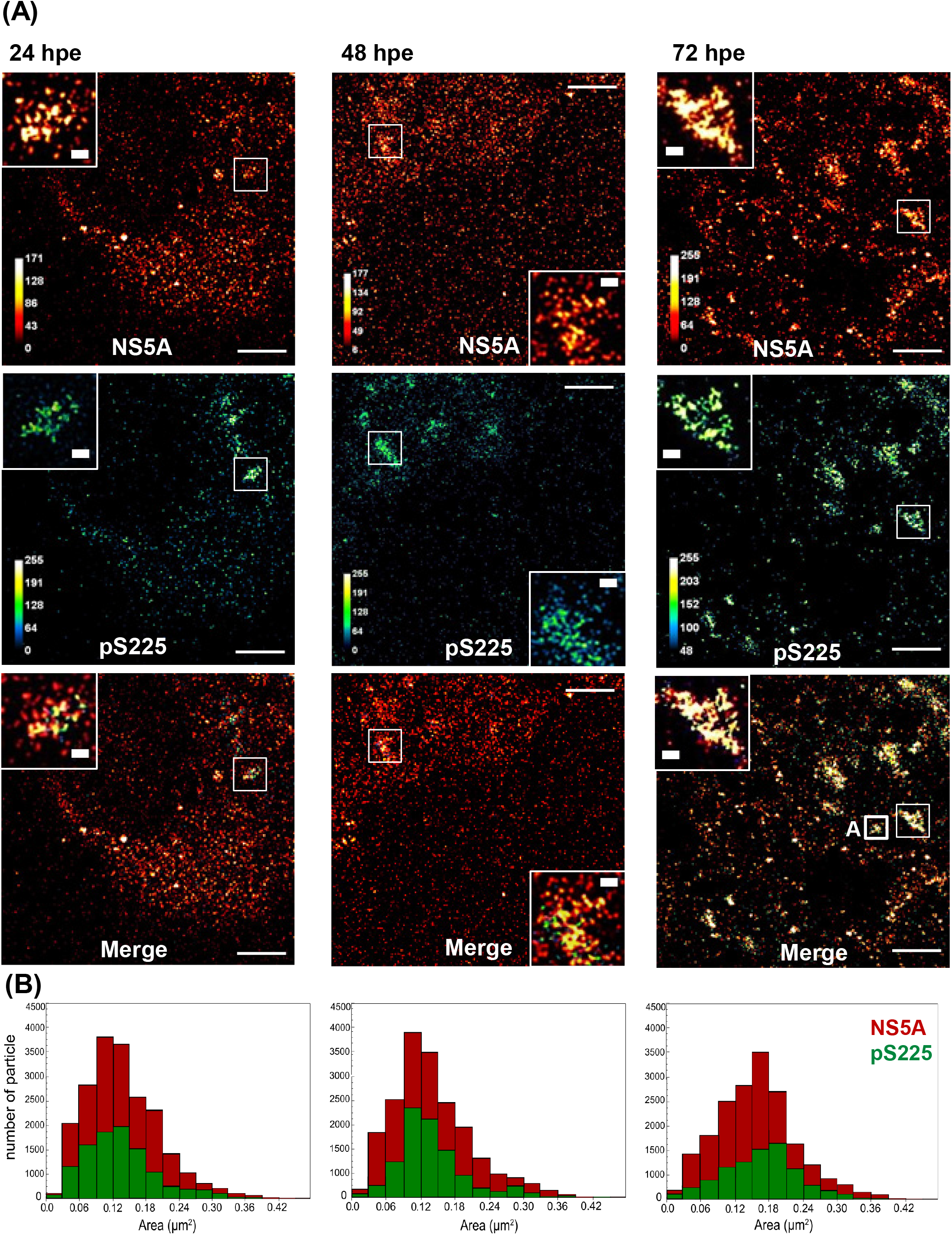
ExM analysis of HCV infected cells. **(A)** Huh7 cells were electroporated with WT mJFH-1 RNA, seeded onto 8-well chambered coverglass and incubated for 24, 48 and 72 h. Cells were fixed and stained for total NS5A (red-hot) and pS225 (green-fire-blue), processed for ExM, and imaged by confocal microscopy. Magnified views of boxed regions and individual channels are shown. Scale bars are 20 μm (physical size post-expansion factor, x4) and 1 μm, respectively. Box **A** corresponds to the cluster that is subject to 3D-reconstruction in Fig 11. **(B)** The size of individual NS5A and pS225-NS5A particles was determined and plotted as a frequency histogram. The area (μm^2^) is taken as an indication of the three-dimensional volume of the NS5A structure.

For comparison, cells harbouring wildtype or S225A mutant SGR-Neo-JFH1 were also analysed by ExM. As expected wildtype NS5A was distributed throughout the cytoplasm (Fig 10A), whereas S225A was restricted to the perinuclear region (Fig 10B). Interestingly, the proportion of NS5A punctae that were pS225 positive was higher than for virus-infected cells (Fig 10B green bars). Wildtype NS5A positive punctae were on average smaller (peak 0.09-0.12 μm^2^) than those in infected cells, however the S225A mutant exhibited significantly condensed and larger punctae (peak 0.18-0.21 μm^2^). It has been reported that treatment of HCV-infected cells with the NS5A inhibitor DCV resulted in a perinuclear accumulation of NS5A (13), reminiscent of that observed for the S225A mutant. We therefore treated wildtype SGR harbouring cells with DCV and analysed the distribution of total NS5A and pS225-NS5A by ExM (Fig 10C). Significant clustering of NS5A punctae was observed and size analysis revealed a bimodal distribution with two populations of punctae exhibiting small (peak 0.06-0.09 μm^2^) and large (peak 0.21 −0.24 μm^2^) areas. The large population corresponded to those punctae observed in S225A mutant SGR harbouring cells.

**Figure 10.**
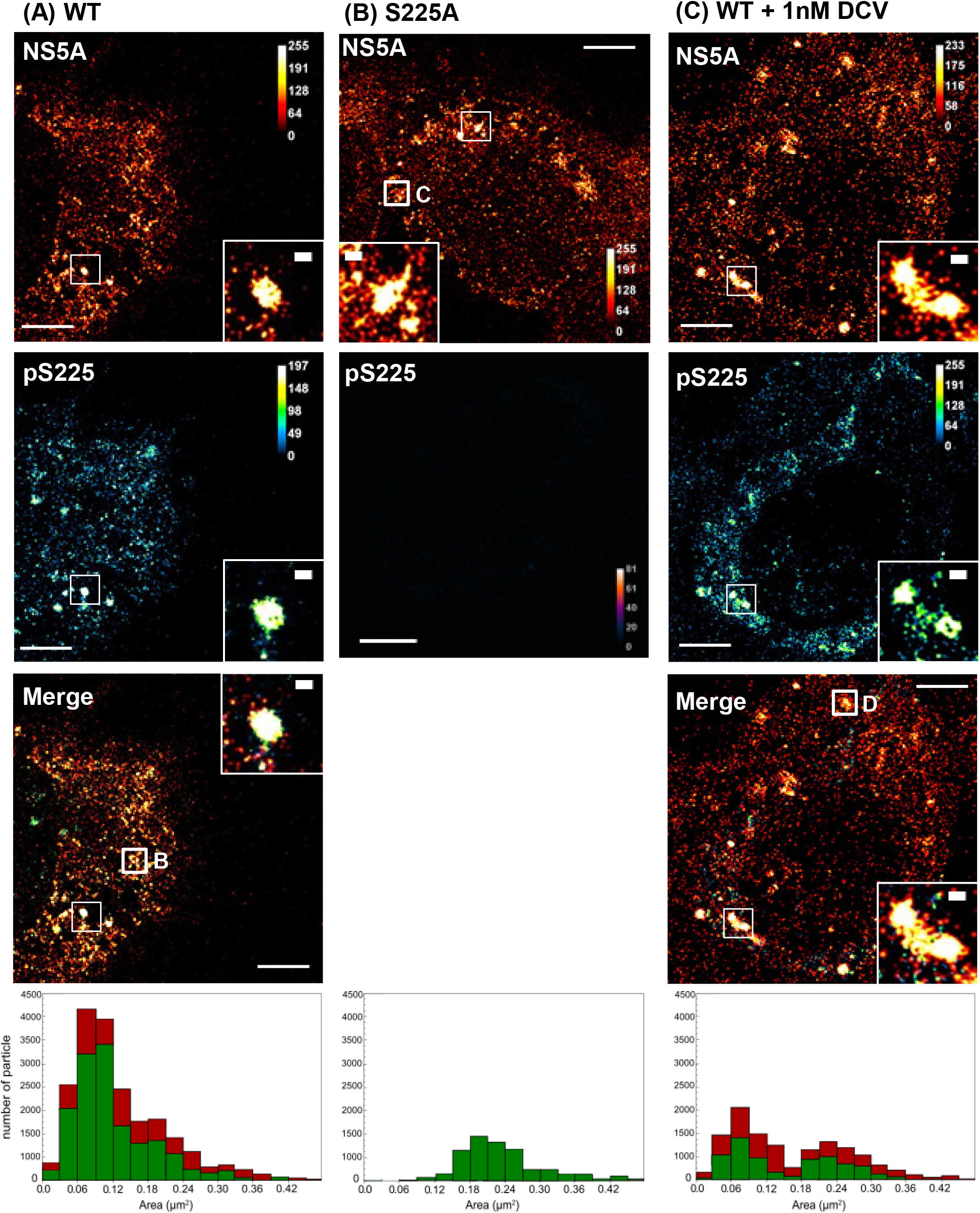
ExM analysis of SGR-harbouring cells. Huh7 cells harbouring either WT or S225A SGR were seeded onto 8-well chambered coverglass and incubated for 48 h. Cells were fixed and stained for total NS5A (red-hot) and pS225 (green-fire-blue), processed for ExM, and imaged by confocal microscopy. Magnified views of boxed regions and individual channels are shown. Scale bars are 20 μm (physical size post-expansion factor, x4) and 1 μm, respectively. (A) WT, (B) S225A and (C) WT following treatment with 1nM DCV for 24 h. Boxes **B, C** and **D** correspond to the clusters that are subject to 3D-reconstruction in Fig 11. The size of individual NS5A and pS225-NS5A particles was determined and plotted as a frequency histogram (lower panels). The area (μm^2^) is taken as an indication of the three-dimensional volume of the NS5A structure.

Next, to fully exploit the improved resolution afforded by ExM, we undertook a 3D-reconstruction of individual clusters of NS5A-positive punctae. Individual clusters from mJFH-1-infected cells (Fig 9A, box A), wildtype SGR-harbouring cells (Fig 10A, box B), S225A SGR-harbouring cells (Fig 10B, box C) or wildtype SGR-harbouring cells treated with DCV (Fig 10C, box D) were chosen for this analysis. In Fig 11 the left panel shows the maximum intensity projections (MIP) of individual clusters that were used to calculate the localisation volumes, followed by 3 different profiles of the clusters (top, bottom and side) (for more examples see Supplementary Fig S3). In mJFH-1 infected cells pS225 was found in distinct areas on the outside surface of the clusters, mostly adjacent to holes that extended through the clusters. The diameter of these holes varied from 0.17-0.4 μm, consistent with LDs. Technical limitations of ExM means that BODIPY (558/568)-C_12_ dye cannot be anchored to the gel matrix and therefore LDs cannot be directly visualised. In SGR-harbouring cells pS225 was more widely distributed over the surface of NS5A clusters, but was still predominantly close to holes. The compact structure of the clusters was not seen in the S225A mutant SGR-harbouring cells, here NS5A occupied larger irregular-shaped volumes and the spaces that likely represent LDs were usually found at the edge. Intriguingly, NS5A clusters in DCV-treated cells exhibited the same expanded, irregular structures as seen for S225A with pS225 distributed across these clusters often at the edges.

**Figure 11.**
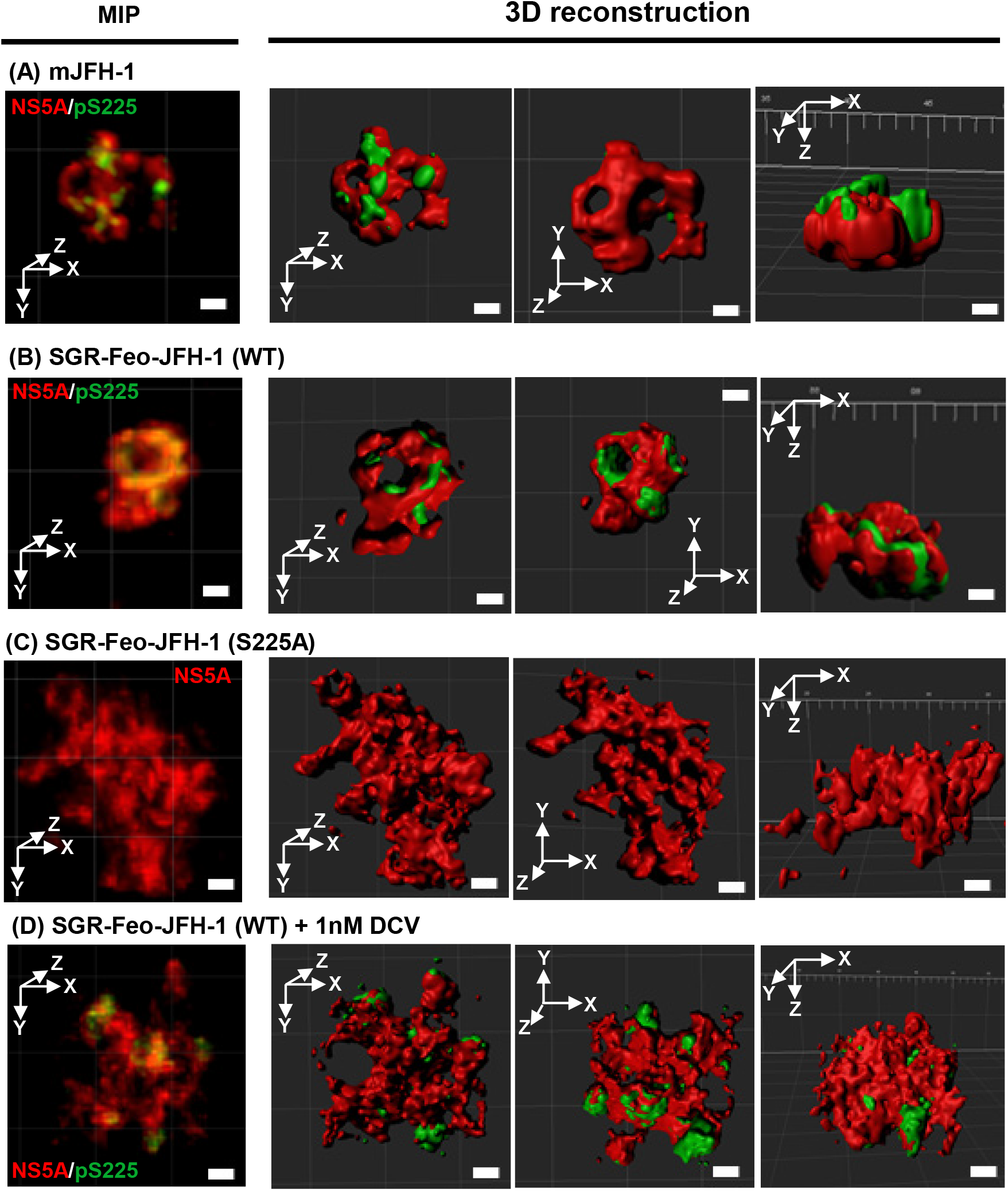
3D reconstruction of clusters of NS5A and pS225-NS5A. The left column shows post-expansion maximum intensity projections (MIP), the remaining columns show 3D reconstructions of NS5A clusters in 3 different orientations (as indicated by X, Y and Z arrows). (A) mJFH-1, (B) SGR-Feo-JFH-1 (WT), (C) SGR-Feo-JFH-1 (S225A) and (D) SGR-Feo-JFH-1 (WT) following treatment with 1nM DCV for 24 h. Scale bars: 1 μm.

To quantify this we measured intensity profiles across the clusters (Fig 12). In both mJFH-1-infected cells and wildtype SGR-harbouring cells intensities measured along vertical and horizontal axis contained two peaks separated by a deep trough, confirming that NS5A was localised around a hole. In contrast the S225A SGR, or wildtype SGR treated with DCV, exhibited more peaks and shallow troughs, suggesting a loss of the tight association with LDs.

**Figure 12.**
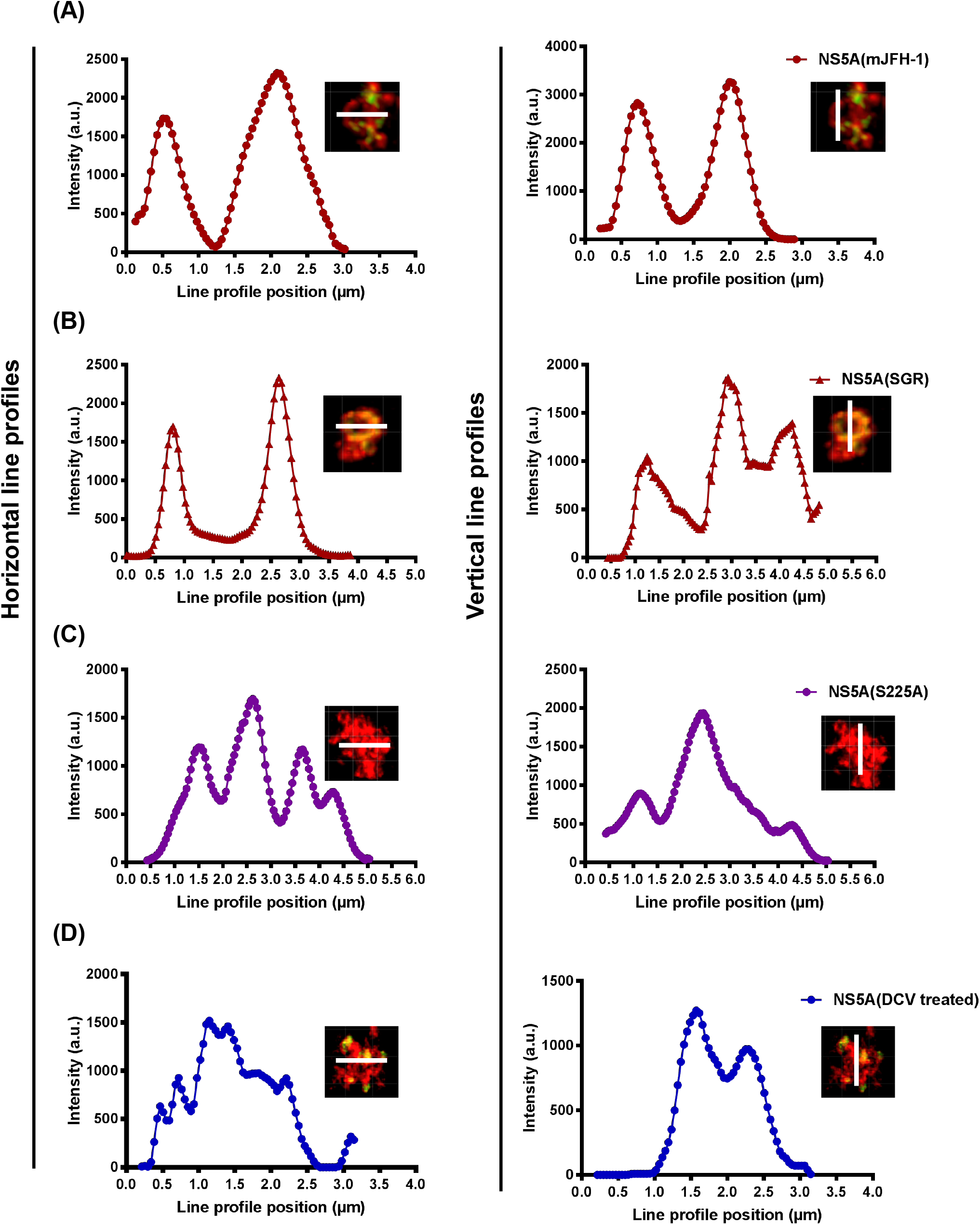
Quantification of ExM maximum intensity projections. Cross-sectional profiles were determined for the four clusters shown in Fig 11. Graphs show line profiles derived from horizontal (left column) and vertical (right column) white lines. (A) mJFH-1, (B) SGR-Feo-JFH-1 (WT), (C) SGR-Feo-JFH-1 (S225A) and (D) SGR-Feo-JFH-1 (WT) following treatment with 1nM DCV for 24 h. All distances and scale bars are in pre-expansion units.

## Discussion

Despite extensive study over many years, the functions of NS5A remain to be unambiguously defined, an issue brought into sharp focus by the advent of DAAs that are predicted to target NS5A. These compounds are highly effective at inhibiting HCV replication, both *in vitro* and in patients, yet virtually nothing is known about their mode of action, or indeed which of the many NS5A functions they inhibit. In this regard, a growing body of evidence points to phosphorylation as a key regulator of NS5A function, however, it is only recently that the phosphorylation sites and kinases that phosphorylate them have begun to be identified. Our studies have focused on S225 in the serine-rich LCSI between DI and DII, for reasons outlined in the introduction. Here we describe the use of an unique reagent – an antiserum specific for pS225 that has allowed us for the first time to directly interrogate the role of S225-phosphorylated NS5A.

Firstly, we showed that pS225 was exclusively present on the p58 NS5A species and identified CKIα and PLK1 as candidate S225 kinases. Pharmacological inhibition of either kinase resulted in a loss of pS225 and p58, but also resulted in a reduction in the overall levels of NS5A. This is likely due to an effect on genome replication and is consistent with the 10-fold reduction in replication seen previously with the S225A phosphoablatant mutant (37). We also observed that inhibition of these kinases resulted in a loss of co-localisation of NS5A with LDs, suggesting that this association is required for efficient genome replication, in addition to its requirement for virus assembly (19, 47).

Previously we had observed that the S225A phosphoablatant mutant had an extensive phenotype (23, 37) and we considered this a disproportionate effect of a single serine-alanine substitution in a highly serine-rich region (Fig 1A). This prompted us to consider that S225 phosphorylation might drive additional proximal phosphorylation events in a sequential (or hierarchical) fashion. Our data are consistent with this (Fig 5) and we propose a model whereby S225 phosphorylation primes subsequent phosphorylation in a bidirectional fashion. Confirmation of this model would require use of additional phospho-specific antibodies, such as those used by Yu *et al* (33) to demonstrate that S238 phosphorylation is dependent on S235 phosphorylation. This observation is consistent with our findings and the model proposed in Fig 13. We propose that phosphorylation of S225 leads to multiple phosphorylation events within a short sequence (17 amino acids), producing a closely packed cluster of phosphates and negative charges with implications for both protein-protein interactions and the conformation of NS5A, for example by changing the orientation of DI and DII respective to each other.

**Figure 13.**
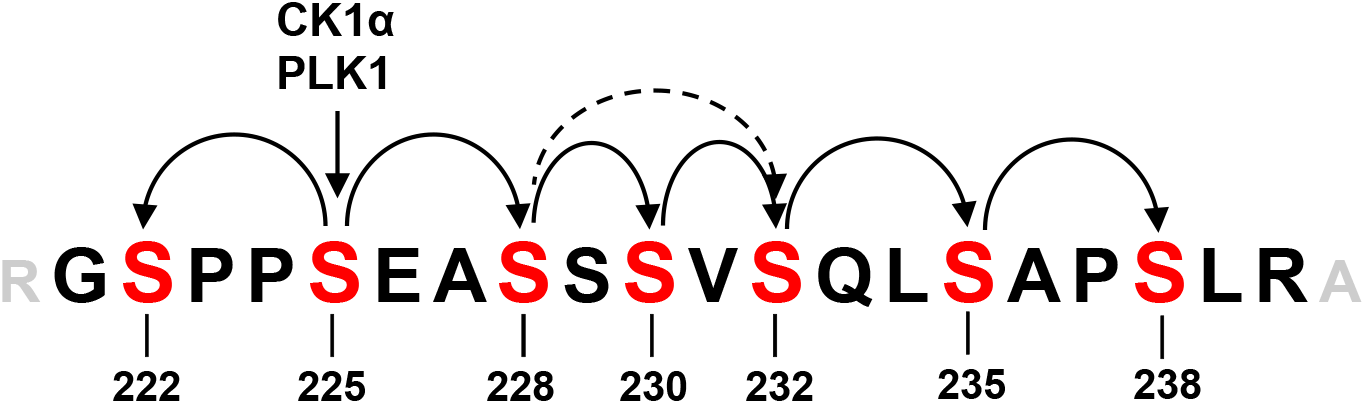
Schematic of hierarchical phosphorylation across LCSI. A proposed model of the location and relationship of phosphorylation events. CK1α and PLK1 phosphorylate S225 and this instigates a bi-directional phosphorylation cascade as shown in curved arrows. The precise details of which serines are phosphorylated between S228-S232 remains obscure as they do not exactly fit to the expected SxxS spacing.

We then combined the pS225 antiserum with two super-resolution microscopy approaches – Airyscan and ExM – techniques which overcome the light diffraction limit of conventional microscopy, to interrogate the subcellular distribution of pS225-NS5A in comparison to the total pool of NS5A. This revealed that pS225-NS5A was only a subset of the total, consistent with the presence of both p56 (lacking pS225) and p58 (pS225-positive) species by western blot (Fig 1B). However, surprisingly we observed that pS225 was not uniformly distributed across the total NS5A (Fig 7). Three-dimensional reconstruction of NS5A clusters imaged by ExM revealed that pS225 foci were surface exposed and in many cases were close to holes that extended through the cluster (Fig 11). These holes are likely to represent LDs which cannot be directly visualised by ExM. These observations lead to a number of speculative conclusions: firstly, the location of pS225 near to LDs is consistent with the loss of NS5A:LD association following treatment with kinase inhibitors and suggests that pS225 is required for the stable interaction of NS5A with LDs. It is conceivable that the concentration of phosphorylated serines in LCSI might mediate interactions with lipids on the surface of LDs. There is precedent for this: for example during lipolysis, cytosol to LD translocation of hormone sensitive lipase (HSL) is driven by phosphorylation of two adjacent serines by protein kinase A, and the LD resident protein perilipin is also extensively phosphorylated during this process (48). Secondly, the surface location of pS225-NS5A would facilitate its interactions with other cellular factors, including cytosolic kinases, explaining the marked effect of S225A (23), or pharmacological inhibition of CKIα and PLK1 (Figs 3, 4), on the distribution of NS5A. ExM also allowed us to observe dramatic alterations in the architecture of NS5A clusters resulting from either the S225A phosphoablatant mutation, or DCV treatment of wildtype (Fig 11). The compact and smooth architecture of the wildtype NS5A cluster with holes (LDs) extending through the structure was replaced with an expanded and irregular structure with less distinct holes. The similarities suggest that S225 phosphorylation and DCV might regulate some common function(s) of NS5A, possibly involved in protein-protein and/or protein-lipid interactions. Again, this is consistent with previous studies showing that DCV treatment disrupted the structure, distribution and formation of membrane-associated replication complexes (49, 50). As a note of caution, it should be noted that these previous studies were performed with NS3-5B expression constructs, rather than in the context of HCV infection.

In conclusion, our study provides more evidence supporting the critical role of phosphorylation in regulating the function of NS5A. In particular we propose that S225 phosphorylation is the primary site within LCSI that primes subsequent phosphorylation events. The high degree of conservation of this residue, and the adjacent serines in LCSI, in all genotypes of HCV suggests some common mechanism across HCV isolates. However, it is intriguing that in the closely related equine hepacivirus (EqHV) (51) the LCSI serine cluster is conserved but the residue corresponding to S225 is aspartate (D). S225D is a phosphomimetic substitution, mimicking the negative charge of the phosphate and possibly maintaining the functionality of pS225. In the future comparative studies between different hepaciviruses may help to elucidate the role(s) of phosphorylation in regulating NS5A function.

## Materials and Methods

### Plasmids

DNA constructs used were derived from the full-length pJFH-1 virus (5) and the luciferase sub-genomic replicon (pSGR-luc-JFH-1) (37, 52). The mSGR-Feo-JFH-1 and mJFH-1 constructs used throughout the study contained unique restriction sites flanking the NS5A coding sequence (20). The serine-alanine mutants have been described previously (37).

### Antibody to pS225

Rabbits were immunised with the 14-mer peptide CARGSPP**pS**EASSS conjugated to KLH. IgG was depleted using the non-phosphorylated peptide and then purified by enrichment using the phospho-peptide (DC Biosciences, Dundee).

### Cell culture

Huh7 cells were maintained in Dulbecco’s modified Eagle’s medium (DMEM; Sigma-Aldrich) supplemented with 10% foetal bovine serum (FBS), 100 IU penicillin/ml, 100 μg streptomycin/ml, and 1% non-essential amino acids (NEAA) in a humidified incubator at 37°C with 5% CO_2_. Huh7 cells carrying a subgenomic JFH-1 replicon (SGR-Feo-JFH-1) were maintained in the same medium supplemented with G418 (300 μg/ml: BioPioneer).

### Electroporation

4×10^6^ Huh7 cells (washed twice and suspended in PBS) were electroporated with 2 μg of RNA at 975 μF and 260 V. Cells were resuspended in complete media and plated in 96-well (3×10^4^cells/well), or 6-well plates (3×10^5^ cells/well).

### Luciferase-based sub-genomic replicon assay

SGR-Feo-JFH-1 harbouring Huh7 cells (2.5×10^4^/well of a 96-well plate), cells were harvested into 30 μl passive lysis buffer (PLB; Promega) per well. Luciferase activity was determined on a BMG plate reader with luciferase assay reagent (Promega).

### Kinase inhibition assays

D4476 (Generon), SBE13-HCl or GSK-3β Inhibitor VIII (Merck) were dissolved in DMSO. Huh7 cells harbouring SGR-Feo-JFH-1 were seeded in a 6-well plate (6×10^5^/well), incubated for 24 h and treated with inhibitors for further 24 h.

### SDS-PAGE/Western blot

Cells were washed twice in PBS, lysed in GLB (1% (v/v) Triton X-100, 120 mM KCl, 30 mM NaCl, 5 mM MgCl2, 10% (v/v) glycerol, 10 mM PIPES-NaOH, pH 7.2 with protease and phosphatase inhibitors) and harvested by centrifugation (2800g, 10 min, 4°C). Following SDS-PAGE, proteins were transferred to a polyvinylidene difluoride (PVDF) membrane and blocked in 50% (v/v) Odyssey blocking (OB) buffer (LI-COR) in TBS. Membranes were incubated with primary antibodies overnight at 4°C, followed by secondary antibodies for 2 h at room temperature (RT), both prepared in 25% OB buffer. Primary antibodies were; anti-NS5A (sheep, 1:4,000) (25), anti-pS225 (1:1000) and mouse α-Actin (Sigma, 1:10,000). Secondary antibodies were anti-rabbit, anti-sheep (800 nm) or anti-mouse (700 nm), used at 1:10,000 prior to imaging using a LI-COR Odyssey Sa infrared imaging system. Quantification was carried out using Image Studio v3.1 (LI-COR) using a background subtraction method.

### Immunofluorescence and confocal microscopy

Cells were washed with PBS, fixed for 20 mins in 4% (w/v) PFA, permeabilised in PBS/0.1% (v/v) Triton X-100 (PBS-T), blocked with PBS-T/5% (w/v) BSA before staining with primary antibody (sheep anti-NS5A 1:2000, rabbit anti-pS225; 1:1000). Fluorescently conjugated secondary antibodies were used at 1:500 (Life Technology). Nuclei were counterstained with DAPI. Lipid droplets were stained with BODIPY (558/568)-C12 (1:1,000, Life Technology). Confocal microscopy images were acquired on a Zeiss LSM880 microscope with Airyscan, post-acquisition analysis was conducted using Zen software (Zen version 2015 black edition 2.3, Zeiss) or Fiji (v1.49) software.

### Expansion microscopy (ExM)

Cells were fixed and stained as above, prior to treatment with 0.1 mg/ml Acryloyl-X, SE (6-((acryloyl)amino) hexanoic acid, succinimidyl ester (AcX) in PBS, (stock 10 mg/ml in DMSO), overnight at RT. For gelation, ammonium persulfate (APS) and tetramethylethylenediamine (TEMED) (0.2% (w/w) each) were added to the monomer solution (1x PBS, 2 M NaCl, 8.625% (w/w) sodium acrylate, 2.5% (w/w) acrylamide, 0.15% (w/w) N,N′- methylenebisacrylamide). The inhibitor 4-hydroxy-2,2,6,6-tetramethylpiperidin-1-oxyl (4-HT) (0.01% (w/w)) was also added. Gelation was performed for 30 min at RT in the same 8 well chamber glass slides, placing a non-adhesive glass slide over the top of the gasket to seal the chamber. Digestion was performed by incubating gels on a rocker overnight at RT with proteinase K (16 units/mL in 50 mM Tris pH 8, 1 mM EDTA, 0.5% Triton X-100, 0.5 M NaCl). Gels were expanded by placing them in excess volumes of double-deionized water for 30 min, replacing the water for additional 30- min washes 4 times.

### Co-localisation analysis

Manders' overlap coefficients were calculated using Fuji ImageJ software with Just Another Co-localisation Plugin (JACoP) (National Institutes of Health) (53). Co-localisation calculations were performed on >5 cells from at least three independent experiments.

## Acknowledgements

We thank Takaji Wakita for the JFH-1 clone and Douglas Ross-Thriepland for the mJFH-1 virus mutants. We also thank Ed Boyden (Massachusetts Institute of Technology) and Isuru Jayasinghe (University of Leeds) for help and advice with expansion microscopy, and Michelle Peckham for access to the Zeiss LSM880 Airyscan confocal microscope (funded by Wellcome Trust grant number 104918/Z/14/Z). We are grateful to Tim Tellinghuisen (Scripps Institute, Florida) for the 9E10 NS5A monoclonal antibody. This work was funded by a Wellcome Trust Investigator Award to MH (grant number 096670/Z/11/Z).

## Author Contributions

Performed experiments: NG CY

Data analysis: NG

Wrote the manuscript: NG MH

Methods: NG

Experimental planning and interpretation: NG MH

Reviewing and editing: NG MH CY

